# Development and characterization of a non-human primate model of disseminated synucleinopathy

**DOI:** 10.1101/2023.12.15.571818

**Authors:** Alberto J. Rico, Almudena Corcho, Julia Chocarro, Goiaz Ariznabarreta, Elvira Roda, Adriana Honrubia, Patricia Arnaiz, José L. Lanciego

## Abstract

The presence of a widespread cortical synucleinopathy is the main neuropathological hallmark underlying clinical entities such as Parkinson’s disease with dementia (PDD) and dementia with Lewy bodies (DLB). There currently is a pressing need for the development of non-human primate (NHPs) models of PDD and DLB to further overcome existing limitations in drug discovery. Here we took advantage of a retrogradely-spreading adeno-associated viral vector serotype 9 coding for the alpha-synuclein A53T mutated gene to induce a widespread synucleinopathy of cortical and subcortical territories innervating the putamen. Four weeks post-AAV deliveries animals were sacrificed and a comprehensive biodistribution study was conducted, comprising the quantification of neurons expressing alpha-synuclein, rostrocaudal distribution and their specific location. In brief, cortical afferent systems were found to be the main contributors to putaminal afferents (superior frontal and precentral gyrus in particular), together with neurons located in the caudal intralaminar nuclei and in the substantia nigra pars compacta (leading to thalamostriatal and nigrostriatal projections, respectively). Obtained data extends current models of synucleinopathies in NHPs, providing a reproducible platform enabling the adequate implementation of end-stage preclinical screening of new drugs targeting alpha-synuclein.

## Introduction

The presence of a widespread synucleinopathy through the cerebral cortex represents the main neuropathological hallmark that typically characterizes Parkinson’s disease with dementia (PDD) and dementia with Lewy bodies (DLB; Martin et al., 2023). Although PDD and DLB overlap in many features, several distinctive characteristics are also observed (Jellinger and Korczyn, 2018), therefore some concerns still remain dealing with to what extent these disorders can be either categorized as two different clinical entities or merely represents the same disease in different stages of progression (Jellinger and Korczyn, 2018; Jellinger, 2022). Among others, levodopa replacement therapies and cholinesterase inhibitors represent the current gold-standard clinical management for PDD and DLB, these pharmacological approaches only leading to a mild symptomatic relief without inducing any disease-modifying effect.

The development of novel therapeutic approaches for Parkinson’s disease (PD) and related synucleinopathies is severely limited by the lack of adequate animal models that properly mimic the known underlying neuropathology (López, 2010; Jansen et al., 2023; Chocarro et al., 2023). Of particular concern, the lack of reliable non-human primate models (NHPs) of PDD and DLB still is an unmet need. Within the field of PD, NHPs are often regarded as the gold-standard model, since several therapeutic interventions such as deep brain stimulation (Rosenow et al., 2004) and repeated apomorphine administration (Luquin et al., 1993) have been made available from research conducted in the MPTP model of PD in NHPs (DeLong, 1990). Although this neurotoxin-based model has been instrumental in setting up most of our current knowledge of basal ganglia function and dysfunction (Lanciego et al., 2012), important limitations apply when considering testing disease-modifying therapeutics and indeed this model failed to recapitulate the neuropathological hallmarks characterizing PD.

Since the use of MPTP-treated NHPs in drug discovery is no longer useful for providing compelling evidence for efficacy and bearing in mind that PD mainly is a synucleinopathy, the field quickly moved towards the implementation of novel NHP models based on the use of adeno-associated viral vectors (AAVs) coding for either wild-type or mutated variants of the alpha-synuclein gene (SNCA, reviewed in Visanji et al., 2016; Koprich et al., 2017; Marmiom and Kordower, 2018). Most of the implemented approaches for AAV-mediated gene transfer of alpha-synuclein (α-Syn) in NHPs were based on direct intranigral parenchymal deliveries in different species, such as marmosets (Kirik et al., 2003; Eslamboli et al., 2007; Bourdenx et al., 2015), rhesus macaques (Yang et al., 2015) and cynomolgus macaques (Koprich et al., 2016; Sucunza et al., 2021). More recent arrivals to the field of capsid engineering enabled the development of several capsid variants leading to a circuit-specific retrograde transduction, these including AAV2-retro (Tervo et al., 2016), AAV-TT (Tordo et al., 2018), AAV-HBKO (Naidoo et al., 2018), AAV-MNM004 and AAV-MNM008 (Davidsson et al., 2019). By taking advantage of retrogradely spreading AAVs, multiple transduction can be achieved in brain areas innervating the injected site, bearing in mind that these capsids behave similarly to traditional retrograde neuroanatomical tracers, at least to some extent (Lanciego and Wouterlood, 2011, 2020). However, it is also worth noting that a reliable retrograde transduction in the NHP brain has been reported for native AAV serotypes such as AAV2, AAV5, AAV6, AAV8 and AAV9 (Markakis et al., 2010; Masamizu et al., 2011; San Sebastian et al., 2013; Gerits et al., 2015; Green et al., 2016).

Among the different AAV serotypes and capsid variants available, AAV9 was here selected according to its accurate and predictable spread through well-characterized brain circuits regardless the strength of these efferent systems, while avoiding non-specific retrograde transduction (Green et al., 2016). Accordingly, an AAV9 coding for the SNCA gene with the A53T mutation under the control of human synapsin promoter [(synapsin)AAV9-SynA53T] was chosen for intraparenchymal deliveries into the left putamen in two cynomolgus macaques (*Macaca fascicularis*) to induce a retrograde transduction of brain areas innervating the putamen in an attempt to generate a widespread synucleinopathy throughout cortical and subcortical brain areas. The conducted biodistribution study was made of three different levels of analysis, comprising (i) a quantification of the total number of transduced neurons within pre-defined regions of interest (ROIs), (ii) rostrocaudal distribution within each ROI and (iii) and accurate location of neurons showing α-Syn expression.

## Materials and Methods

### Study design

This study was aimed to develop and characterize a NHP model of disseminated synucleinopathy mimicking the known neuropathological signatures of PDD and DLB to the best possible extent. Accordingly, we sought to determine whether the intraputaminal delivery of a retrogradely spreading AAV9 coding for mutated alpha-synuclein (AAV9-SynA53T) is able to induce a widespread synucleinopathy throughout cortical and subcortical brain areas innervating the putamen. Two adult juvenile NHPs (*Macaca fascicularis*) were injected with AAV9-SynA53T into the left putamen and sacrificed four weeks post-AAV deliveries. Upon animal sacrifices, brain tissue samples were processed for histological analysis and up to three different readouts were considered, comprising number of transduced neurons, rostrocaudal distribution and location.

### Experimental animals

A total of two adult juvenile naïve *Macaca fascicularis* NHPs (34 months-old; both females; body weight between 2.35 and 2.49 Kg) were used in this study. Animal handling was conducted in accordance to the European Council Directive 2010/63/UE as well as in full keeping with the Spanish legislation (RD53/2013). The experimental design was approved by the Ethical Committee for Animal Testing of the University of Navarra (ref: CEEA095/21) as well as by the Department of Animal Welfare of the Government of Navarra (ref: 222E/2021).

### Viral vector production

A recombinant AAV vector serotype 2/9 expressing the SNCA gene with A53T mutation driven by the human synapsin promoter was produced by Vector Builder (https://en.vectorbuilder.com/; catalog No. VB201126-1235dnc). The virus was formulated in PBS buffer (pH 7.4) supplemented with 200 nM NaCl and 0.001% pluronic F-68. Obtained vector concentration was 9.62 x 10^13^ GC/ml. Virus purity was determined by SDS-PAGE followed by silver staining, resulting in >80% pure. Plasmid map for pAAV-synapsin-SynA53T and sequence are provided in Supplementary Figure 1.

### Stereotaxic surgery for AAV deliveries

Surgical anesthesia was induced by intramuscular injection of ketamine (5 mg/Kg) and midazolam (0.5 mg/Kg). Local anesthesia was implemented just before the surgery with a 10% solution of lidocaine. Analgesia was achieved with a single intramuscular injection of flunixin meglumine (Finadyne®, 5 mg/Kg) delivered at the end of the surgical procedure and repeated 24 and 48 h post-surgery. A similar schedule was conducted for antibiotic coverage (ampicillin, 0.5 ml/day). After surgery, animals were kept under constant monitoring in individual cages with ad libitum access to food and water. Once animals showed a complete post-surgical recovery (24 h), they were returned to the animal vivarium and housed in groups.

Stereotaxic coordinates for AAV deliveries into the left putamen were calculated from the atlas of Lanciego and Vázquez (2012). During surgery, target selection was assisted by ventriculography. Pressure deliveries of AAVs were made through a Hamilton® syringe in pulses of 5 μl/min for a total volume of 25 μl each into three sites in the left putamen, each deposit spaced 1 mm in the rostrocaudal direction to obtain the highest possible transduction extent of the putamen. Once injections were completed, the needle was left in place for an additional time of 10 min before withdrawal to minimize AAV reflux through the injection tract. Coordinates for the more rostral deposits in the left putamen of AAV9-SynA53T were 1 mm rostral to the anterior commissure (ac), 1 mm dorsal to the bi-commissural plane (ac-pc plane) and 11 mm lateral to the midline. Second deposits were performed at the ac level (ac = 0 mm), 1 mm dorsal to the ac-pc plane and 11.5 mm lateral to the midline, whereas the more caudal injections were conducted 1 mm caudal to ac, 1.5 mm dorsal to the ac-pc plane and 12.5 mm lateral to the midline.

### Necropsy, tissue processing and data analysis

Anesthesia was firstly induced with an intramuscular injection of ketamine (10 mg/Kg), followed by a terminal overdose of sodium pentobarbital (200 mg/Kg) and perfused transcardially with an infusion pump. Both animals (MF04 and MF05) were sacrificed four weeks post-AAV deliveries. The perfusates consisted of a saline Ringer solution followed by 3,000 ml of a fixative solution made of 4% paraformaldehyde and 0.1% glutaraldehyde in 0.125 M phosphate buffer (PB) pH 7.4. Perfusion was continued with 1,000 ml of a cryoprotectant solution containing 10% glycerine and 1% dimethylsulphoxide (DMSO) in 0.125 M PB pH 7.4. Once perfusion was completed, the skull was opened and the brain removed and stored for 48 h in a cryoprotectant solution containing 20% glycerin and 2% DMSO in 0.125 M PB pH 7.4. Next, frozen coronal sections (40 μm-thick) were obtained on a sliding microtome and collected in 0.125 M PB pH 7.4 as 10 series of adjacent sections. These series were used for (1) immunoperoxidase detection of α-Syn, (2) immunoperoxidase detection of α-Syn counterstained with toluidine blue (Nissl stain), (3) immunoperoxidase detection of tyrosine hydroxylase (TH), and (4) immunoperoxidase detection of TH counterstained with toluidine blue. The remaining 6 series of sections were stored at -80 °C cryopreserved in 20% glycerin and 2% DMSO until further use, if needed. A completed list of primary and bridge antisera (secondary biotinylated antisera), together with incubation concentrations, incubation times and commercial sources is provided below in Table 1.

**Table 1:**
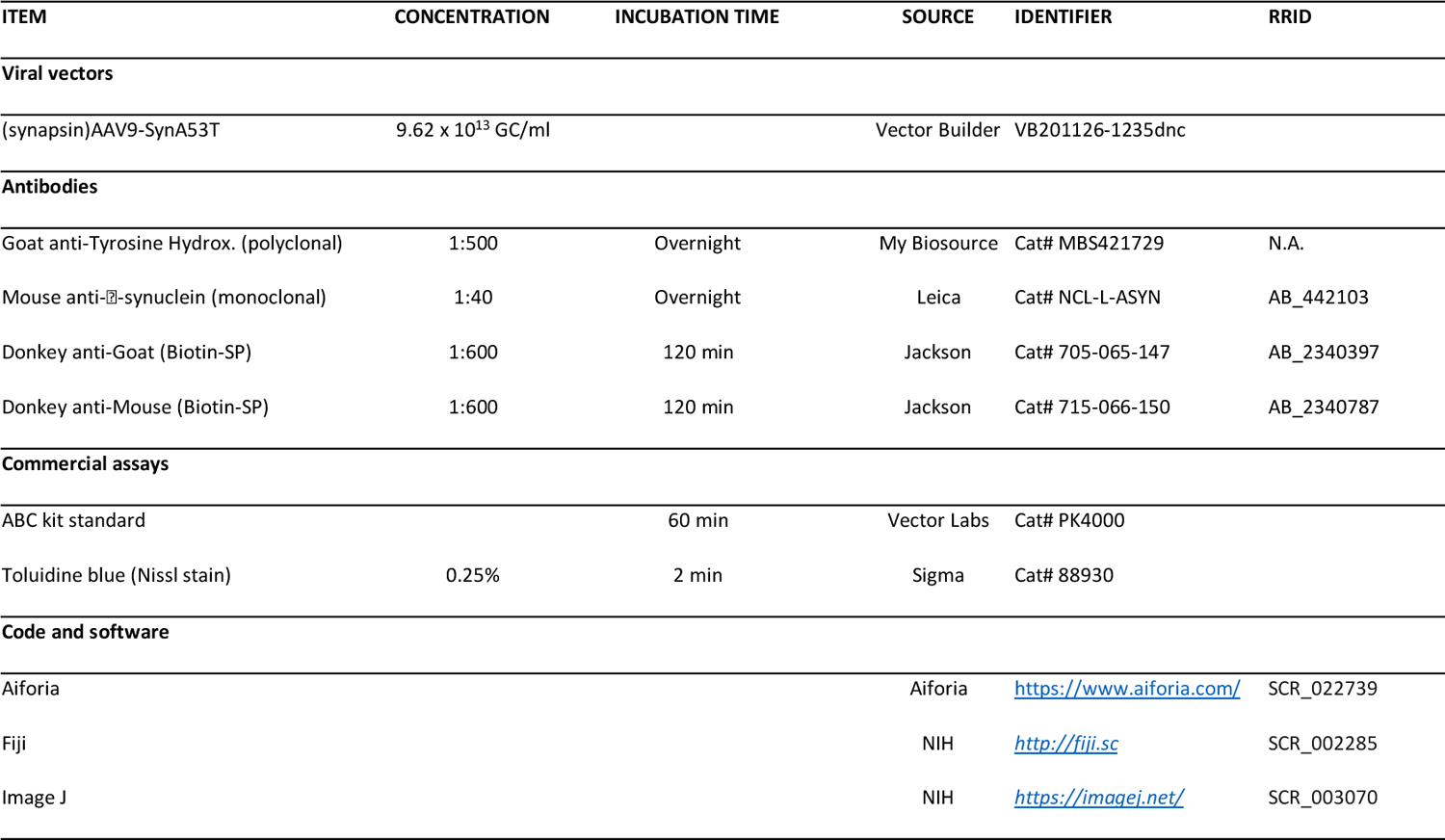
List of reagents & resources.

### Quantification of neurons expressing a-Syn

Every tenth section was processed for the immunoperoxidase detection of α-Syn and used for estimating the number of transduced neurons with AAV9-SynA53T. For this purpose, a deep-learning dedicated algorithm was prepared with Aiforia® (Penttinen et al., 2018; https://www.aiforia.com/), validated and released (resulting in an error of 1.67% for quantifying α-Syn+ neurons). Sixty-nine equally spaced coronal sections covering the range of ac +10 mm to ac -14 mm were sampled per animal, including all ROIs containing α-Syn+ neurons at the cortical (left and right cerebral cortices) and subcortical (left hemisphere) levels. Sections were scanned at 20x in an Aperio CS2 slide scanner (Leica, Wetzlar, Germany). Selected cortical ROIs comprised the anterior cingulate gyrus (ACgG), superior frontal gyrus (SFG), middle frontal gyrus (MFG), inferior frontal gyrus (IFG), precentral gyrus (PrG), fronto-orbital gyrus (FoG), medial orbital gyrus (MOrG), lateral orbital gyrus (LOrG) posterior cingulate gyrus (PCgG), postcentral gyrus (PoG), insular gyrus (Ins), superior parietal lobule (SPL), supramarginal gyrus (SMG), precuneus (PCu), superior temporal gyrus (STG) and middle temporal gyrus (MTG), whereas subcortical ROIs included the amygdaloid complex (AMG), ventral anterior thalamic nucleus (VAL), ventral lateral thalamic nucleus (VL), ventral posterior thalamic nucleus (VPO), centromedian-parafascicular thalamic complex (CM-Pf), substantia nigra pars compacta (SNpc) and dorsal raphe nucleus (DRN). Segmentation of cortical areas was conducted in keeping with the stereotaxic atlas of Martin and Bowden (1996). In brief, scanned sections were uploaded to the Aiforia® cloud. Next, the boundaries of each selected ROI were outlined at low magnification and the dedicated algorithm was then used as a template quantifying the desired neuronal population (e.g. α-Syn+ neurons). Representative images illustrating the accuracy of the conducted procedure are provided in Figure 1.

**Figure 1:**
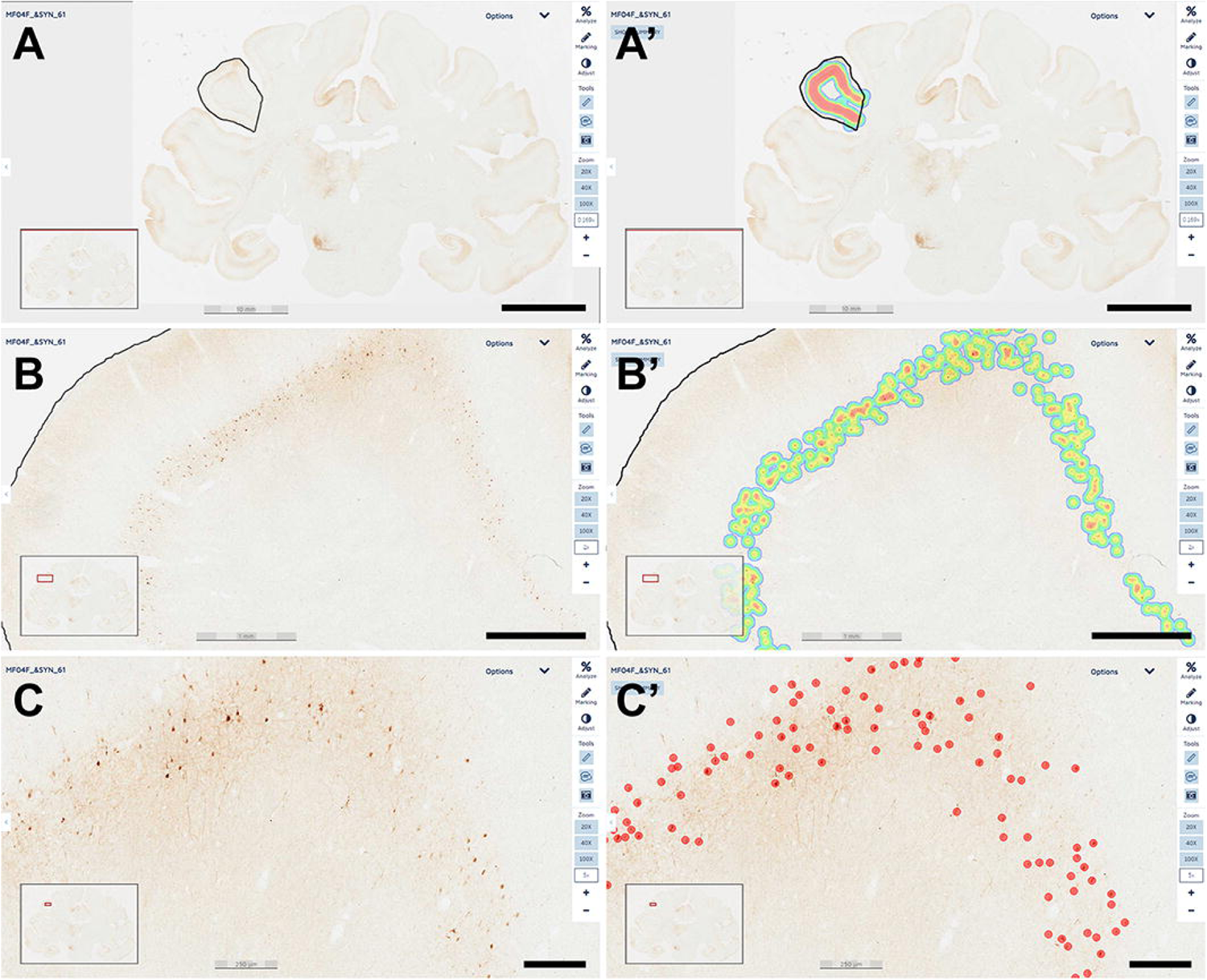
Quantification of α-Syn+ neurons. A dedicated deep-learning algorithm identifying α-Syn transduced neurons was prepared with Aiforia® (www.aiforia.com). Scanned sections were uploaded to Aiforia® cloud and several different ROIs were outlined throughout cortical and subcortical territories. Panel A illustrates the delineation of the left supramarginal gyrus at low magnification (0.169x). Labeling intensity, reflecting the number of transduced neurons is represented color-coded in panel A’. Panel B shows the same area at a magnification of 2x, together with the accompanying result, color-coded as illustrated in panel B’. A higher-magnification view (5x) of the obtained labeling is shown in panel C, together with the accurate identification of every single α-Syn+ neuron, as observed in panel C’. All transduced neurons were quantified within the selected ROI. Illustrated images are taken from animal MF04 and correspond to coronal section #61. In summary, all α-Syn+ neurons were counted, and indeed the algorithm was fully accurate and specific (a total of 414 neurons were identified in this ROI). Scale bars are 10 mm for panels A & A’; 1 mm in panels B & B’ and 250 μm in panels C & C’.

### Quantification of the nigrostriatal system

At the level of the substantia nigra pars compacta (SNpc), the number of TH+ neurons were quantified with a different Aiforia® algorithm designed to disclose TH+ cells (resulting in an error of 4.82%). In both animals, the analysis was conducted in thirteen TH-stained, equally-spaced coronal sections covering the whole rostrocaudal extent of the SNpc. Sections were scanned at 20x in a slide scanner (Aperio CS2; Leica), and uploaded to the Aiforia® cloud (https://www.aiforia.com/) where TH+ neurons were quantified according to a similar procedure described above for estimating the number of α-Syn+ neurons. Regarding the density of TH+ terminals at the level of the putamen, up to 26 equally-spaced coronal sections stained for TH and covering the whole rostrocaudal extent of the left and right putamen (pre-and post-commissural) in each animal were scanned at 20x and used for measuring TH optical densities with Fiji Image J (NIH, USA; https://imagej.net/software/fiji/) and coverted to a logarithmic scale according to available protocol (Ruifrok and Johnston, 2011).

## Results

A basal ganglia circuit-specific widespread synucleinopathy was generated upon the delivery of a retrogradely-spreading AAV9-SynA53T. The conducted study enabled the design of a novel NHP model of disseminated synucleiniopathy mimicking the known neuropathology of PDD and DLB, at least to some extent. Detailed biodistribution patterns for α-Syn+ neurons innervating the putamen through cortical and subcortical locations are reported here.

### Injection sites and transduced putaminal areas with AAV9-SynA53T

The delivery of AAV9-SynA53T in animals MF04 and MF05 (3 sites, 25 μl each of the viral suspension) were accurate and properly located within the boundaries of the putamen, showing a minimal spread of the viral suspension towards neighboring white matter tracts surrounding the putamen and without any noticeable reflux through the injection tract (Figure 2A & B). In both animals, a broad transduced area in the putamen was observed, ranging from 64.97% to 67.18% (animals MF04 and MF05, respectively) of the total rostrocaudal extent of the injected putamen (Figures 2C, 2D and 2E).

**Figure 2:**
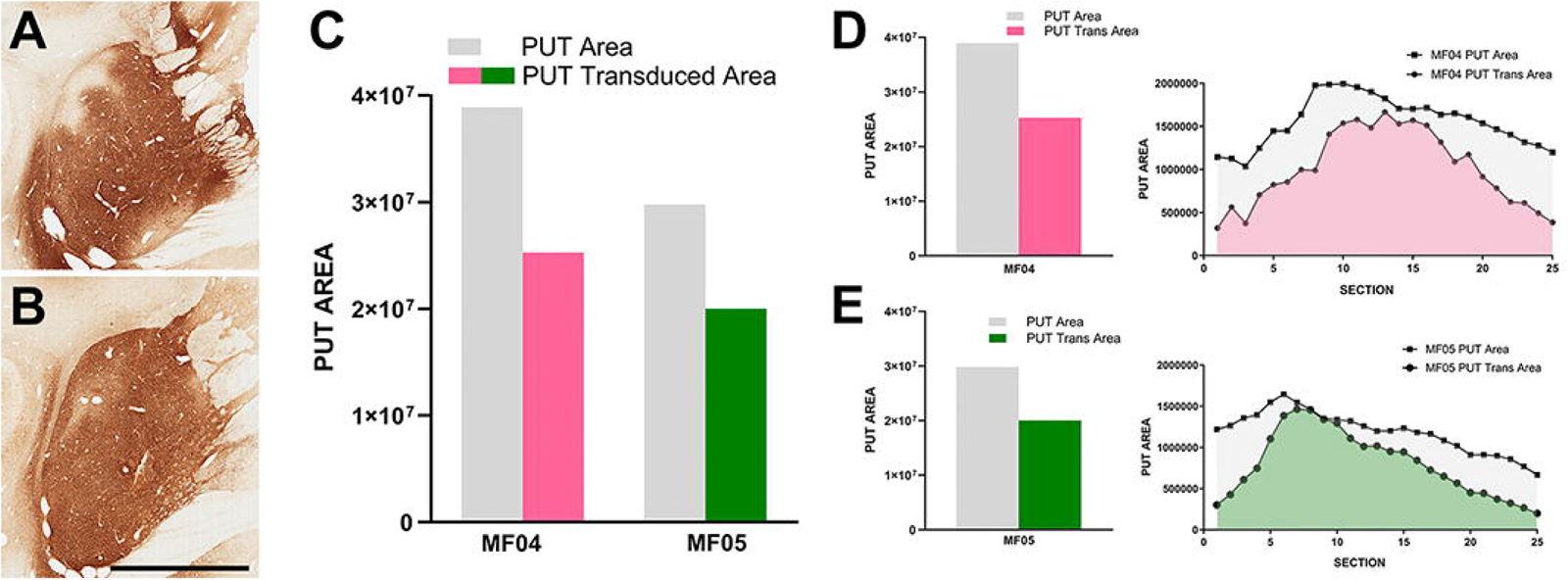
Injection sites and transduced putaminal areas. Panels A & B illustrate the intraparenchymal deliveries of AAV9-SynA53T in the putamen at the level of ac = 0 mm (animal MF04 is represented in panel A; animal MF05 is illustrated in panel B). The extent of the putaminal transduced area is shown in the histogram of panel C. The area of the left putamen is represented as grey bars, whereas the extent of transduced areas are illustrated as either purple (animal MF04) or green bars (animal MF05). Panels D & E represent the rostrocaudal extent of the transduced area (purple for animal MF04 and green for animal MF04) against the rostrocaudal extent of the putamen (grey). Upon the delivery of AAV9-SynA53T into the putamen (3 sites; 25 μl each), obtained transduced areas accounted for 64.97% and 67.18% (animals MF04 and MF05, respectively) of the total putaminal area. Viral vector expression was largely restricted within putaminal boundaries, with minimal spread towards neighboring white matter tracts surrounding the putamen. Furthermore, a complete lack of viral spread through the injection tracts can easily be observed in panels A & B (scale bar = 7.0 mm).

### Number of a-Syn+ neurons across pre-defined ROIs

Transduced neurons were located in cortical and subcortical structures innervating the putamen through well-characterized circuits (Lanciego et al., 2012), and the total number of α-Syn+ neurons is reflecting the different strengths of striatal afferent systems.

Considering cortical ROIs, the precentral and superior frontal gyri were the territories containing the highest number of α-Syn+ neurons, followed by the anterior and posterior cingulate gyri. A lower number of neurons was found within the postcentral, insular and supramarginal gyri. Although consistent in both animals, a sparser labeling was observed in the middle and inferior frontal gyri, orbital cortex (lateral and medial orbital territories), superior and middle temporal gyri, superior parietal lobule and precuneus. As expected, cortical areas engaged in motor processing such as the primary motor area and the supplementary motor area (located within the precentral gyrus and superior frontal gyrus, respectively) are those showing the highest number of α-Syn+ transduced neurons (Figure 3). Regarding contralateral cortico-putaminal projections, the highest number of α-Syn+ neurons was always found in the precentral and superior frontal gyri, with a more moderate labeling being observed in the anterior and posterior cingulate cortices. Fewer transduced neurons were found in contralateral territories comprising the middle and inferior frontal gyri, the supramarginal gyrus, the postcentral gyrus and the superior parietal lobule. Finally, α-Syn+ neurons were never found in contralateral cortices such as the lateral and medial orbital gyri (Figure 3).

**Figure 3:**
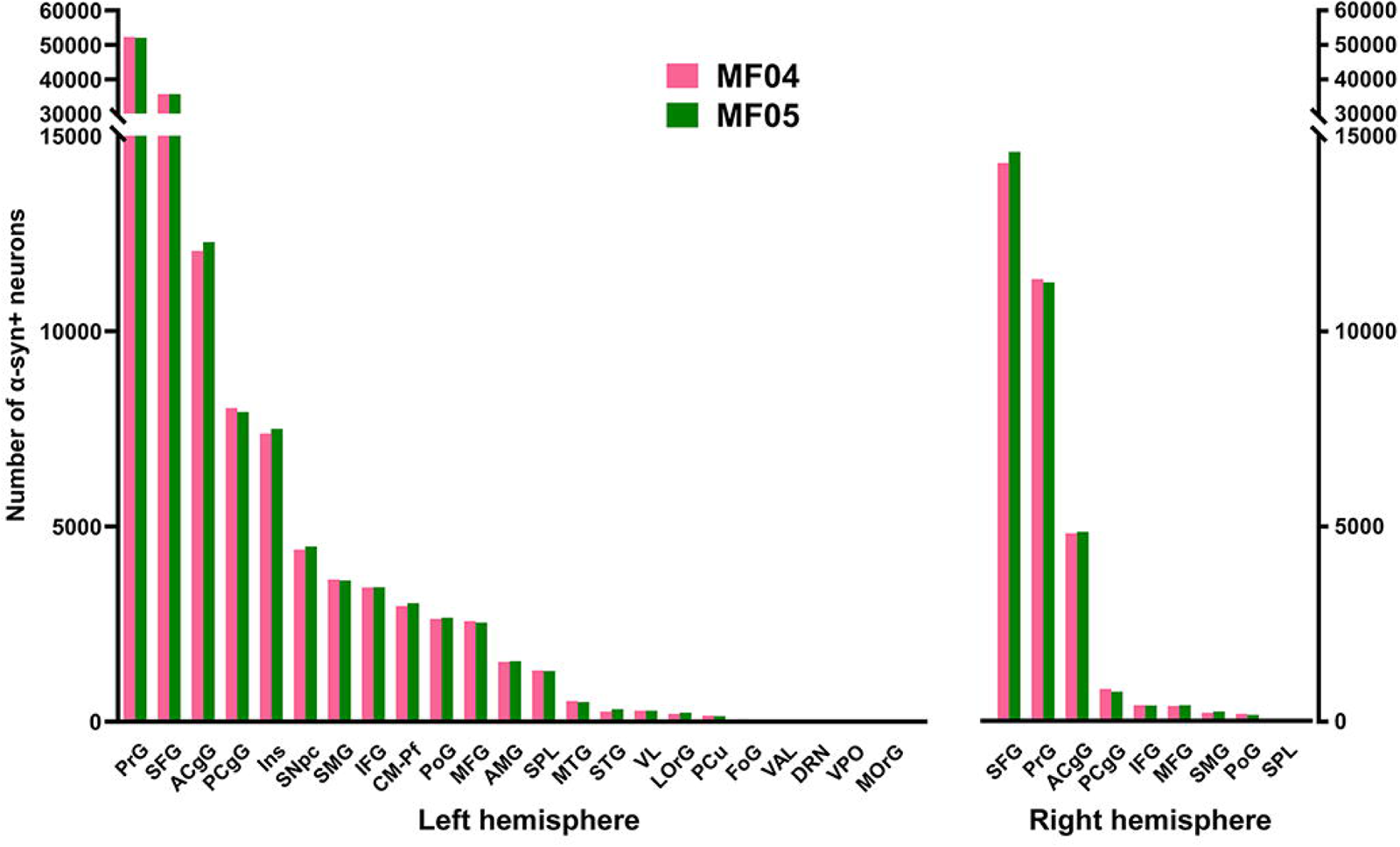
Number of α-Syn+ neurons across pre-defined ROIs. Histograms showing the total number of quantified α-Syn+ neurons within each selected ROI in the left and right hemispheres. Both animals showed very similar numbers of transduced neurons within each territory. The prefrontal and superior frontal gyri are by large the main contributors to corticostriatal projections (left and right hemispheres), followed by the cingulate gyri (anterior and posterior; both hemispheres), as well as the insular gyrus (left hemisphere only). At the subcortical level, the caudal intralaminar nuclei and the substantia nigra pars compacta of the left hemisphere are the areas containing the higher number of a-Syn+ neurons. Abbreviations (from left to right in histograms): precentral gyrus (PrG), superior frontal gyrus (SFG), anterior cingulate gyrus (ACgG), posterior cingulate gyrus (PCgG), insular gyrus (Ins), substantia nigra pars compacta (SNpc), supramarginal gyrus (SMG), inferior frontal gyrus (IFG), centromedian-parafascicular thalamic complex (CM-Pf), postcentral gyrus (PoG), middle frontal gyrus (MFG), amygdaloid complex (AMG), superior parietal lobule (SPL), middle temporal gyrus (MTG), superior temporal gyrus (STG), ventral lateral thalamic nucleus (VL), lateral orbital gyrus (LOrG), precuneus (PCu), fronto-orbital gyrus (FoG), ventral anterior thalamic nucleus (VAL), dorsal raphe nucleus (DRN), ventral posterior thalamic nucleus (VPO), and medial orbital gyrus (MOrG).

At subcortical levels, the highest number of α-Syn+ neurons was found in the substantia nigra pars compacta and the centromedian-parafascicular thalamic complex, with a smaller participation of the amygdaloid complex, in keeping with the expected strength of nigrostriatal, thalamostriatal and amygdaloid-striatal unilateral efferent systems (Figure 3).

Although initially not designed for this purpose, the conducted quantification of α-Syn+ neurons enabled the estimation of the strength of each individual striatal afferent system, namely corticostriatal (ipsi-and contralateral), thalamostriatal and nigrostriatal projections. Ipsilateral corticostriatal projections are by far the most abundant ones, representing on average up to 72.13% of the total number of α-Syn+ neurons. Within this projection system, main contributors to striatal afferents are those arising from the superior frontal and precentral gyri that collectively accounted for 70.74% of the total ipsilateral corticostriatal projections. Contralateral corticostriatal projection neurons comprised 18.95% of the total striatal afferents (78.82% of these neurons being located in the contralateral superior frontal and precentral gyri). When compared with corticostriatal projections, striatal afferents arising from subcortical territories like the centromedian-parafascicular thalamic complex and the substantia nigra pars compacta accounted for much lower percentages (1.73% and 2.58%, respectively). It is worth noting that contralateral corticostriatal projections (18.95% of the total number of α-Syn+ neurons) represent a percentage relevant enough for a projection that has often been neglected in studies dealing with basal ganglia function and dysfunction (Borra et al., 2022).

### Rostrocaudal distribution of a-Syn+ neurons within pre-defined ROIs

Besides quantifying the total number of transduced neurons within any given territory, another parameter that needs to be taken into consideration is the rostrocaudal extent of obtained labeling in each analyzed ROI (e.g. to evaluate how much of the extent for a given ROI contains α-Syn+ neurons). Obtained data revealed that α-Syn+ neurons covered most of the rostrocaudal extent of the anterior and posterior cingulate gyri, the superior, middle and inferior frontal gyri, the precentral gyrus, the postcentral gyrus and the insular gyrus in the left hemisphere. The same also applies to the left centromedian-parafascicular thalamic complex and the left substantia nigra pars compacta. A mirror-like, similar pattern of rostrocaudal distribution is observed in the contralateral hemisphere at the level of the anterior and posterior cingulate gyri, superior frontal gyrus, precentral gyrus and postcentral gyrus. A schematic representation of rostrocaudal biodistribution patterns is provided in Figure 4.

**Figure 4.**
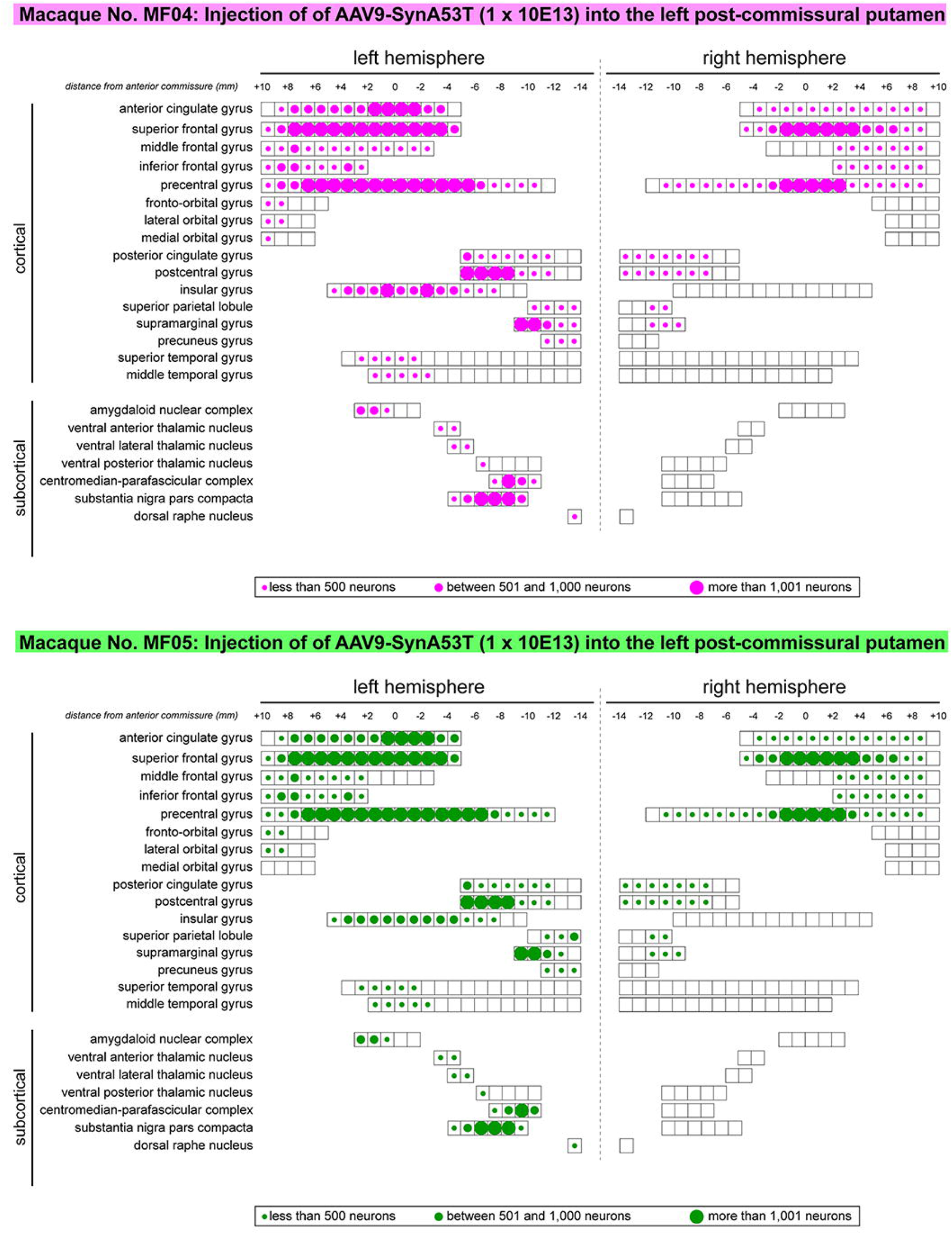
Rostrocaudal distribution of α-Syn+ neurons across pre-defined ROIs. Schematic representation of the rostrocaudal distribution of transduced neurons within each selected cortical and subcortical territories. Several cortical areas contained α-Syn+ neurons throughout their full rostrocaudal extent, as observed in the cingulate gyri (anterior and posterior), superior, middle and inferior frontal gyri, precentral and postcentral gyri, the insular gyrus, as well as in subcortical territories such as the centromedian-parafascicular thalamic complex and the substantia nigra pars compacta. Besides representing the rostrocaudal distribution, dots of different sizes were here used to illustrate the relative number of transduced neurons. Obtained results were very similar in animals MF04 and MF05, with minimal variations.

### Location of a-Syn+ neurons

Addressing the location of transduced neurons represents another important level of analysis, to properly demonstrate the accuracy of circuit-specific retrograde transduction mediated by AAV9-SynA53T. This is best exemplified by the fact that that all neurons expressing α-Syn are located in predictable brain areas, i.e. within brain territories such as the ipsi-and contralateral cortex, ipsilateral amygdaloid complex, caudal intralaminar nuclei and substantia nigra pars compacta, that all together are well-known sources of striatal afferent systems. Moreover, not a single α-Syn+ neuron was found in any given brain area not innervating the putamen. Furthermore, at the level of the cerebral cortex (ipsi- and contralateral cortices), most of the neurons expressing α-Syn were found in deep cortical layers (layer V in particular), where most of the corticostriatal-projecting neurons are known to be located. Only very sparse and inconsistent labeling was found in cortical layers other than layer V. A schematic representation of the location of transduced neurons is provided in Figure 5.

**Figure 5.**
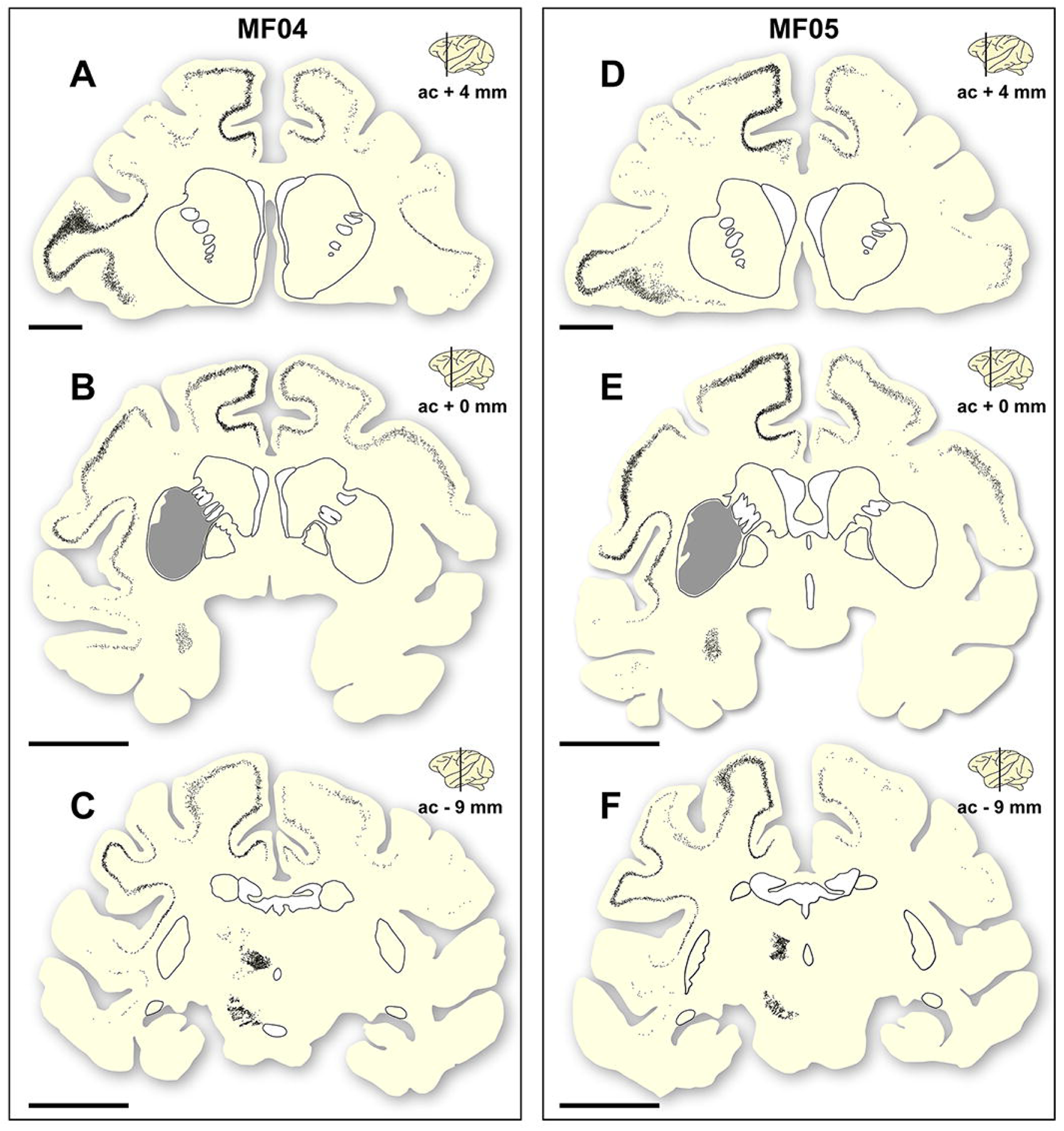
Location of a-Syn+ neurons. Schematic representation of the accurate location of α-Syn+ neurons in the two animals of the study (panels A, B & C are gathered from animal MF04; panels D, E & F belong to animal MF05). Within each animal, up to three coronal levels are illustrated (ac +4 mm; ac +0 mm and ac -9 mm). The location of the caudate and putamen nuclei were delineated for reference purposes. Each dot corresponds to a single α-Syn+ neuron. Furthermore, the extent of the transduced area at ac level of +0 mm was also highlighted as a grey area (panels B & E). At the cortical level, most of the α-Syn+ neurons were located in deep cortical layers (layer V), with only very few neurons being found in cortical layers other than layer V. Furthermore, the density of α-Syn+ neurons at the level of the caudal intralaminar nuclei and the substantia nigra pars compacta are high enough to elucidate the boundaries of these territories. Scale bar is 5 mm in panels A & D, and 10 mm in panels B, C, E & F.

Finally and when considering cortical neurons expressing α-Syn, the intensity of the obtained labeling (e.g. levels of α-Syn expression) ranged from moderate to high across all analyzed ROIs (Figures 6 and 7). Indeed, obtained retrograde labeling with AAV9-SynA53T was sometimes found to be of “Golgi-like” morphology, i.e. not only restricted to the cell somata but also extending to distal dendrites, as seen in several locations throughout the cerebral cortex (Figure 7H).

**Figure 6.**
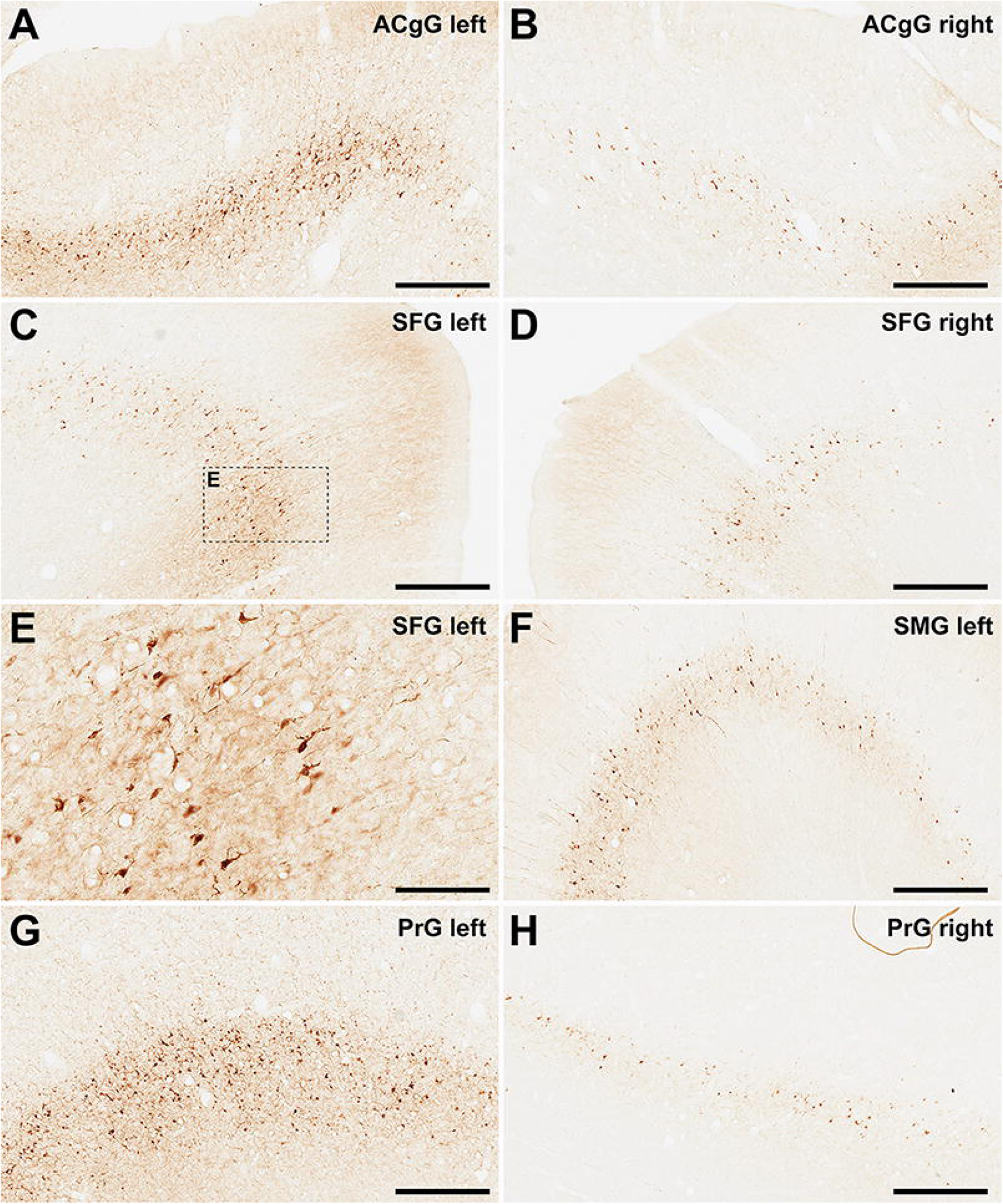
α-Syn+ neurons in animal MF04. Photomicrographs showing the readout of the conducted immunoperoxidase stains with antibodies against alpha-synuclein. Images were taken from the left and right hemispheres to better illustrate observed inter-hemispheric differences in terms of total number of α-Syn+ neurons (anterior cingulate gyrus, superior frontal gyrus, supramarginal gyrus and precentral gyrus). According to conducted quantification, ipsilateral corticostriatal projections accounted for 68.70% of the total number of projection neurons, whereas the contralateral corticostriatal projections represented up to 18.91% of the total striatal afferent systems. Scale bar is 125 μm in panel E and 500 μm in all the remaining panels.

**Figure 7.**
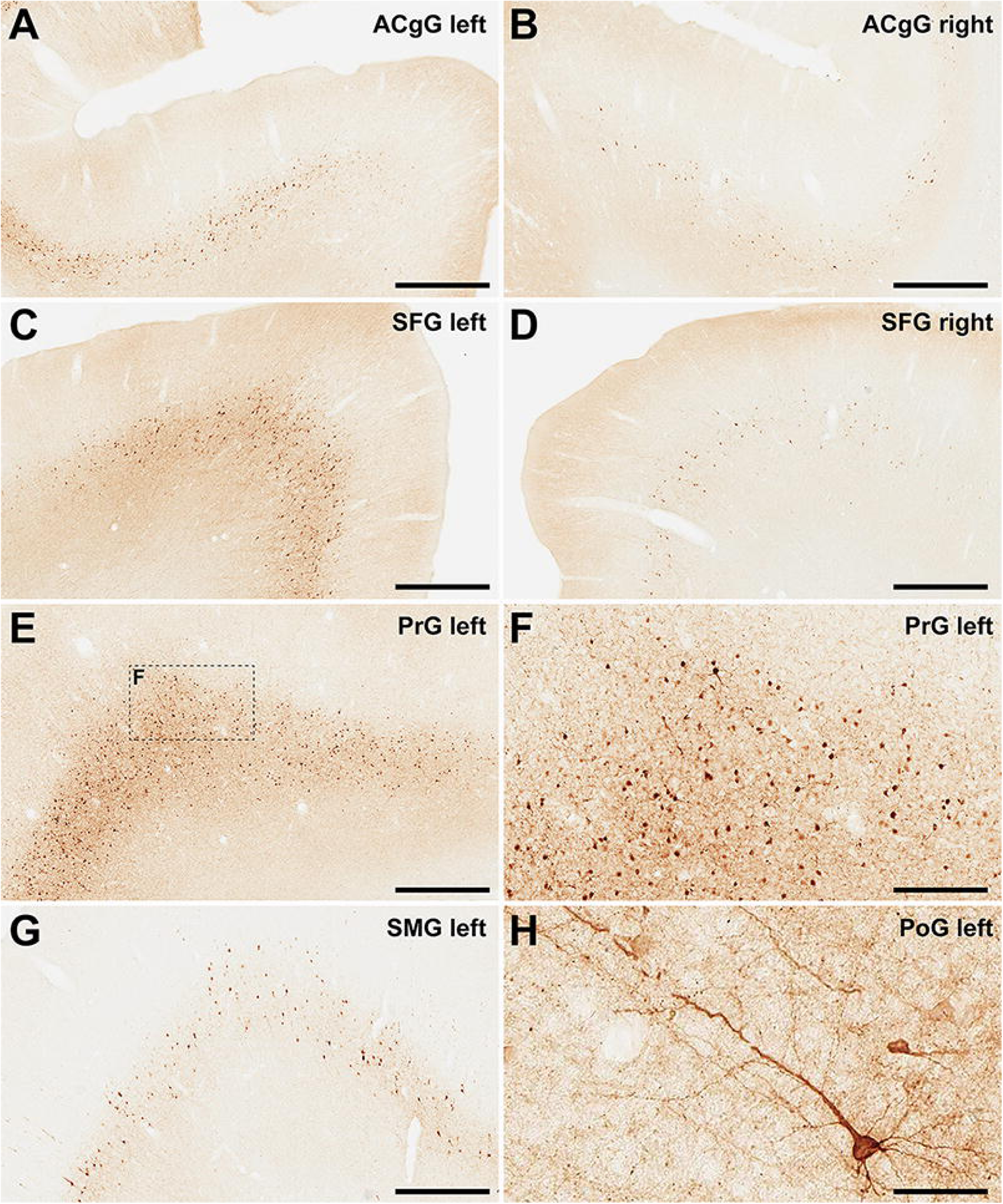
α-Syn+ neurons in animal MF05. Photomicrographs showing the readout of the conducted immunoperoxidase stains with antibodies against alpha-synuclein. Images were taken from the left and right hemispheres to better illustrate observed inter-hemispheric differences in terms of total number of α-Syn+ neurons (anterior cingulate gyrus, superior frontal gyrus, supramarginal gyrus, precentral gyrus and postcentral gyrus). At high magnification, Golgi-like labeling of projection neurons can be sometimes observed, with α-Syn expression comprising the cell somata, basal and apical dendrites (panel H) According to conducted quantification, ipsilateral corticostriatal projections accounted for 75.55% of the total number of projection neurons, whereas the contralateral corticostriatal projections represented up to 18.99% of the total striatal afferent systems. Scale bar is 250 μm in panel F, 62.5 μm in panel H and 500 μm in all the remaining panels.

### Subcortical labeling

The caudal intralaminar nuclei and the substantia nigra pars compacta are known to be the main sources of thalamostriatal and nigrostriatal projections, respectively. Therefore, these two subcortical structures were the ones that concentrated most of the transduced neurons with α-Syn in subcortical territories. Obtained labeling at these levels was abundant enough to allow a proper definition of their boundaries (Figure 5). Sparser labeling was also found in the amygdaloid complex and in the dorsal raphe nucleus, two additional sources of weaker putaminal afferents (Figures 2-5).

### Preserved nigrostriatal system

For any study intended to induce a widespread synucleinopathy in NHPs, additional attention should be placed in analyzing the potential impact of α-Syn expression within nigrostriatal projections, both at origin (e.g. number of TH+ neurons in the substantia nigra pars compacta) as well at destination (e.g. density of TH+ terminals in the putamen). In the left substantia nigra pars compacta, the ratios between number of α-Syn+ neurons and TH+ neurons were similar in both animals (54.49% in MF04 and 56.26% in MF05). Although roughly half of the dopaminergic neurons became transduced with AAV9-SynA53T, the obtained quantification of TH+ neurons in the left and right substantia nigra revealed the lack of dopaminergic cell degeneration in both animals. The same applies to the analysis of optical densities for TH+ terminals in the left and right putamen (Figure 8). In other words, although transduction levels were substantial enough, nigrostriatal damage was not observed.

**Figure 8.**
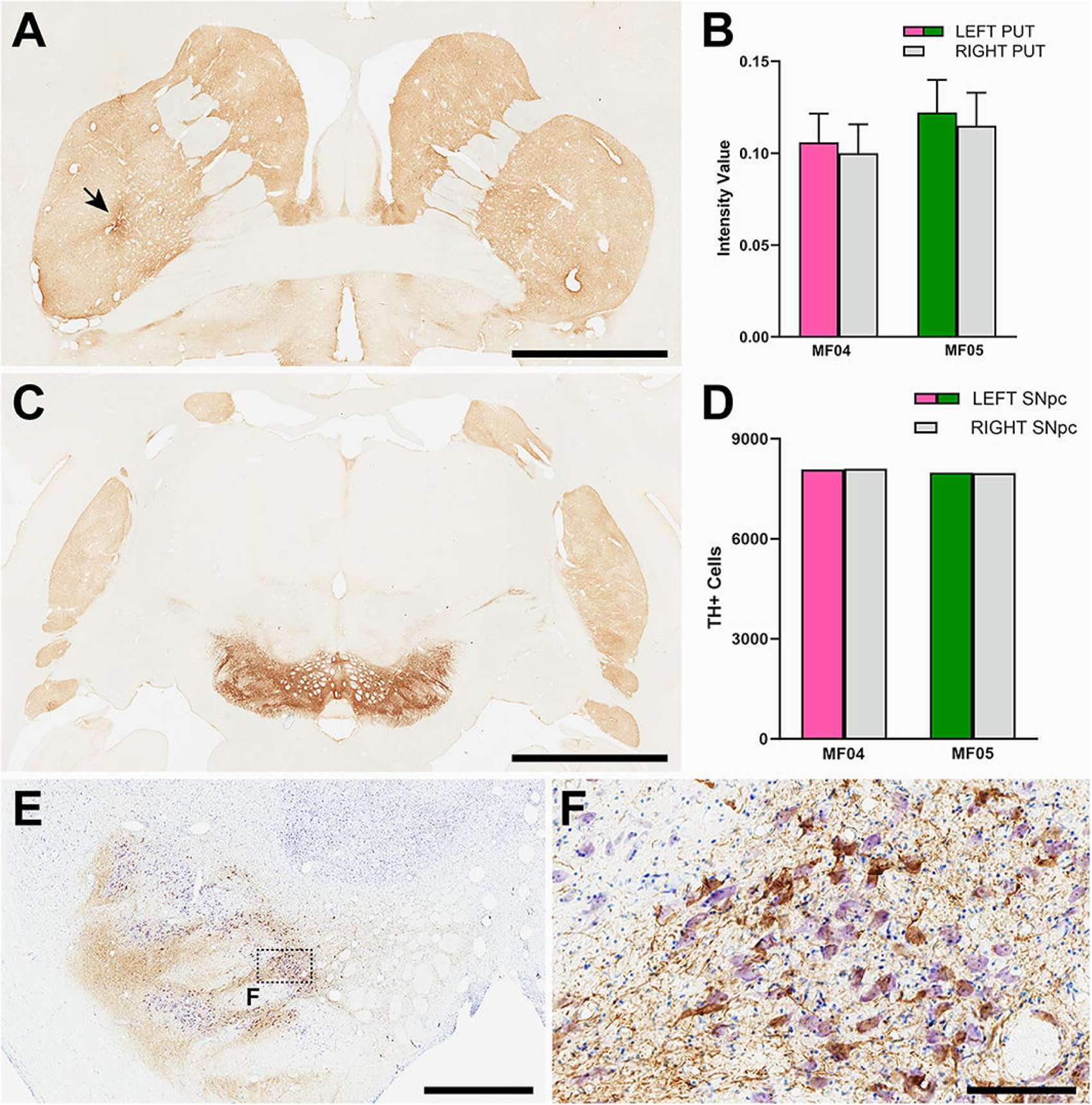
Nigrostriatal system: Photomicrographs and histograms illustrating the conducted analyses at the level of the nigrostriatal system. The optical density of TH+ terminals was compared between the left and right putamen across 26 equally-spaced sections. Panel A shows a coronal section immunostained for TH at the level of the midline decussation of the anterior commissure (animal MF05). Arrow in panel A indicates the injection site of AAV9-SynA53T. Panel B is a histogram showing obtained quantifications for optical densities in both animals. The right hemisphere is represented by a grey bar, whereas purple and green bars represent the left putamen (animals MF04 and MF05, respectively). Optical densities of TH+ terminals were similar when comparing the left and right putamen nuclei in both animals. Panel C is a low-power photomicrograph taken from a coronal section through the substantia nigra in animal MF05, immunostained for TH. As indicated in the histogram shown in panel D, the number of TH+ neurons remain the same when comparing left vs. right substantia nigra (color-coded the same was as in panel B). Panel E and inset (panel F) illustrates the immunoperoxidase detection of α-Syn counterstained with toluidine blue (Nissl stain) in a coronal section through the substantia nigra in animal MF04. Although roughly half of the dopaminergic neurons became transduced with α-Syn (54.94% in MF04 and 56.26% in animal MF05), any evident sign of nigrostriatal damage was observed. Scale bar is 6.5 mm in panel A, 3.5 mm in panel B, 1,750 μm in panel E and 175 μm in panel F.

## Discussion

In this study, the performance and biodistribution of the retrogradely-spreading AAV9-SynA53T vector was evaluated in the NHP brain. Conducted intraparenchymal deliveries of viral suspensions in the left putamen gave rise to a disseminated synucleinopathy in a circuit-specific basis.

### Technical considerations

#### Notes on the chosen nomenclature

Cortical and subcortical ROIs were defined here in keeping with the parcellation conducted in the stereotaxic atlases of Martin and Bowden (1996) and Lanciego and Vázquez (2012) when considering cortical and subcortical structures, respectively. In both references, segmentation was based on anatomical landmarks, such as cortical gyri and sulcus. However, within few other available atlases of the NHP brain (Saleem and Logothetis, 2007; Paxinos et al., 2009), parcellation was based on functional territories, i.e. referring to primary motor and sensory cortices, supplementary motor area, premotor area, etc. In our view, making reference to anatomical landmarks is more accurate and facilitates inter-species comparisons, whereas a precise limitation of functional territories often is more troublesome.

#### Notes on the study design

The relatively low number of animals used in this study (n = 2) may be viewed as a potential limitation for the conducted study, however reflecting the current worldwide shortage of NHP supplies (Janssen et al., 2023). In this context, the need for reduction should also be taken into consideration (e.g. using the lowest possible number of animals) and indeed it is worth noting that obtained results were fully comparable in both animals, with minimal -if any-differences in terms of total number of neurons, rostrocaudal distribution and location of transduced cells.

### Retrogradely-spreading AAV capsid variants

The field of AAV capsid design and engineering is constantly evolving at a breath-taking speed, with new arrivals being made available and tested in a wide range of different animal species, including NHPs (Campos et al., 2023). Pathway-specific retrograde transduction of neurons innervating any given AAV-injected CNS territory collectively represents an appealing scenario for both basic neuroscience as well as for therapeutic interventions. While several different retrogradely-spreading AAV capsids are currently available, selection of the most appropriate one for a given study still represents a difficult choice, which is further confounded by the choice of surrogate animal model for clinical species. An ideal retrograde-propagating AAV capsid should fulfill at least two golden rules, namely (i) accurate and predictable spread through well-characterized brain circuits, further enabling the retrograde transduction of all neurons innervating the injected structure regardless the strength of these efferent systems, while avoiding non-specific retrograde transduction, the latter represented by either AAV uptake through fibers of passage or by the presence of transduced neurons in CNS territories not connected with the injection site and (ii) comparative inter-species performance, with similar transduction efficiency regardless the chosen experimental animal. Furthermore, emphasis should be placed of stereotaxic expertise, with injection sites and transduced areas remaining restricted within the boundaries of the targeted structures without spreading beyond to either neighboring structures or surrounding white matter fiber tracts. In summary, same considerations overall than those applying to stereotaxic deliveries of more traditional retrograde neuroanatomical tracers (Lanciego and Wouterlood, 2011, 2020).

From the available technical arsenal, the AAV capsid variant firstly introduced for retrograde spread was the so-called AAV2-retro (Tervo et al., 2016). Although this capsid exhibits a good performance for the transduction of corticostriatal projections in both rodents and NHPs, its ability to transduce subcortical striatal afferent systems (thalamostriatal and nigrostriatal) is more questionable and indeed notable inter- species differences were reported for AAV2-retro (Cushnie et al., 2020; Weiss et al., 2020; Albaugh et al., 2020). A potential replacement for AAV2-retro is AAV2-HBKO, a viral capsid obtained through mutation of the heparin sulfate proteoglycan binding pocket in AAV2 (Naidoo et al., 2018). Upon MRI-guided deliveries of AAV2-HBKO into the thalamus of NHPs, retrogradely-transduced neurons were found in several territories of the cerebral cortex as well as in subcortical locations such as the caudate and putamen nuclei, globus pallidus and substantia nigra pars reticulata and compacta (Naidoo et al., 2018). Besides AAV2-retro and AAV2-HBKO, several novel AAV capsid variants enabling retrograde transduction have been made available recently, these comprising AAV-MNM004 and AAV-MNM008 (Davidsson et al., 2019), AAV-TT (Tordo et al., 2018) and AAV2i8 (Kondratov et al., 2021). According to available data, AAV2i8 performs nicely in NHPs, whereas AAV-MNM004, AAV-MNM008 and AAV-TT have not been tested yet in NHPs to the very best of our knowledge.

That said and given the well-known good performance of the native AAV9 capsid for retrograde spread through brain circuits (Green et al., 2016; Sucunza et al., 2021), AAV9 still is regarded as a very good alternative to engineered capsids and indeed often represents the standard choice for CNS applications.

### Comparisons with existing AAV-based models of synucleinopathy in NHPs

Upon the identification that alpha-synuclein is the main component of Lewy bodies in idiopathic cases of PD and DLB (Spillantini et al., 1997), the field of PD modeling in NHPs quickly moved to approaches based on the use of either AAVs coding for different forms of the SNCA gene or preformed, synthetic alpha-synuclein fibrils (reviewed in Koprich et al., 2017; Marmion and Kordower, 2018). Initial studies were performed in marmosets by Kirik et al. (2003) by performing intranigral AAVs deliveries coding for either mutated or wild-type forms of alpha-synuclein. After a follow-up period of 16 weeks, a significant reduction in the density of TH+ striatal terminals was noticed, together with 30-60% of TH+ cells in the substantia nigra. Later on, roughly similar results were reported by Eslamboli et al. (2007) in marmosets after intranigral deliveries of AAV2/5 (wild-type and mutated alpha-synuclein) and with a longer follow-up period ranging from 33 to 52 weeks. More recently, another study in marmosets (Bourdenx et al., 2015) used an AAV9 coding for mutated A53T synuclein, injected in the substantia nigra and comprising both young and aged marmosets. Obtained results revealed lower percentages of TH+ cell loss, ranging from 13% in young animals to 20% in aged animals. Regarding macaques, unilateral deliveries in the substantia nigra with a lentiviral vector coding for mutated A53T synuclein in *Macaca mulatta* revealed non-quantified levels of TH cell loss in the substantia nigra (Yang et al., 2015). More recently, the intranigral delivery of AAV1/2 in *Macaca fasicularis* resulted in 50% of TH+ cell loss 17 weeks after AAV deliveries (Koprich et al., 2016). Next and in our own experience, deliveries of AAV9-SynA53T in the substantia nigra of *Macaca fascicularis* lead to 55% of dopaminergic cell loss 12 weeks after viral injections (Sucunza et al., 2021). Finally, a different approach was recently implemented in *Macaca fascicularis* upon the intranigral delivery of an AAV coding for the human tyrosinase gene (the rate-limiting enzyme for neuromelanin sysnthesis). This strategy lead to a time-dependent enhanced accumulation of neuromelanin, later triggering an endogenous synucleinopathy in the form of Lewy bodies (Chocarro et al., 2023).

Obtained results here did not observed any noticeable dopaminergic cell loss in the substantia nigra, and the same holds true when analyzing optical densities of TH+ terminals in the putamen. When compared to existing NHP models of synucleinopathy based on AAVs, two main considerations may arise: firstly, intranigral and intraputaminal deliveries of viral vectors can hardly be compared together, bearing in mind that intranigral deliveries maximize transgene concentration into the targeted area, whereas a retrograde spread from the putamen towards the substantia nigra is required for achieving dopaminergic cell transduction. Secondly and most importantly, huge differences apply when comparing the range of survival times. Sacrifices were performed here four weeks upon intraputaminal deliveries, whereas much longer survival times were used in studies directly targeting the substantia nigra. In other words, although 4 weeks post-delivery of AAV9-SynA53T is an interval good enough to induce α-Syn transduction, longer follow-up times will be likely required to further induce neurodegeneration. Moreover, different AAV serotypes, viral titration and inter- species differences may also account for the observed differences. However, it is worth noting that the obtained transduction of TH+ cells upon intraputaminal deliveries has its own added value, since by going this way the identity of α-Syn+ neurons in the substantia nigra can be disclosed (e.g. transduced neurons are all of them projecting to the putamen), a fact that cannot be properly addressed upon intranigral deliveries (e.g. projection patterns for transduced neurons remained unknown).

## Conclusion

Here a novel NHP model of disseminated synucleinopathy was introduced by taking advantage of the retrograde spread of AAV9-SynA53T upon intraparenchymal putaminal deliveries. This model likely represents a good standard for the pre-clinical development of new drugs intended to induce alpha-synclein clearance for the treatment of unmet medical conditions such as PDD and DLB.

## Supporting information

plasmid map (pAAV-CMV-hTyr) & sequence

## Data availability

Further information and request for resources and reagents should be directed to and will be fulfilled by the corresponding authors, José L. Lanciego.

Full datasets can be found at Zenodo repository: https://doi.org/10.5281/zenodo.10390251

Detailed descriptions of the conducted protocols is available at protocols.io repository:

http://dx.doi.org/10.17504/protocols.io.e6nvwd3w7lmk/v1

http://dx.doi.org/10.17504/protocols.io.rm7vzxzw5gx1/v1

https://www.protocols.io/view/protocol-for-34-immunoperoxidase-detection-of-alph-c9kvz4w6

https://www.protocols.io/view/protocol-for-34-quantification-of-neurons-expressi-c9nsz5ee

https://www.protocols.io/view/protocol-for-34-quantification-of-the-nigrostriata-c9nrz5d6

## Ethics statement

Animal handling was conducted in accordance to the European Council Directive 2010/63/UE as well as in full keeping with the Spanish legislation (RD53/2013). The experimental design was approved by the Ethical Committee for Animal Testing of the University of Navarra (ref: CEEA095/21) as well as by the Department of Animal Welfare of the Government of Navarra (ref: 222E/2021).

## Author contributions

AJR and JLL designed the experiments and performed NHP surgeries and sacrifices with the assistance of JC and GA. AC, JC and GA collected data and performed data analysis with feedback from ER, AH and PA. AJR, AC and JLL wrote and revised the manuscript. All authors contributed to the article and approved the submitted version.

