## Supplementary material for "Development and characterization of a non-human primate model of disseminated synucleinopathy": plasmid map (pAAV-CMV-hTyr) & sequence

### Vector Summary

| | |
|---|---|
| Vector ID | VB201126-1235dnc |
| Vector Name | pAAV[Exp]-SYN1>{hSNCA[NM_001375286.1]*(A53T)}:WPRE |
| Vector Size | 4701 bp |
| Viral Genome Size | 2104 bp |
| Vector Type | Mammalian Gene Expression AAV Vector |
| Inserted Promoter | SYN1 |
| Inserted ORF | {hSNCA[NM_001375286.1]*(A53T)} |
| Inserted Regulatory Element | WPRE |
| Plasmid Copy Number | High |
| Antibiotic Resistance | Ampicillin |
| Cloning Host | VB UltraStable (or alternative strain) |

### Vector Map

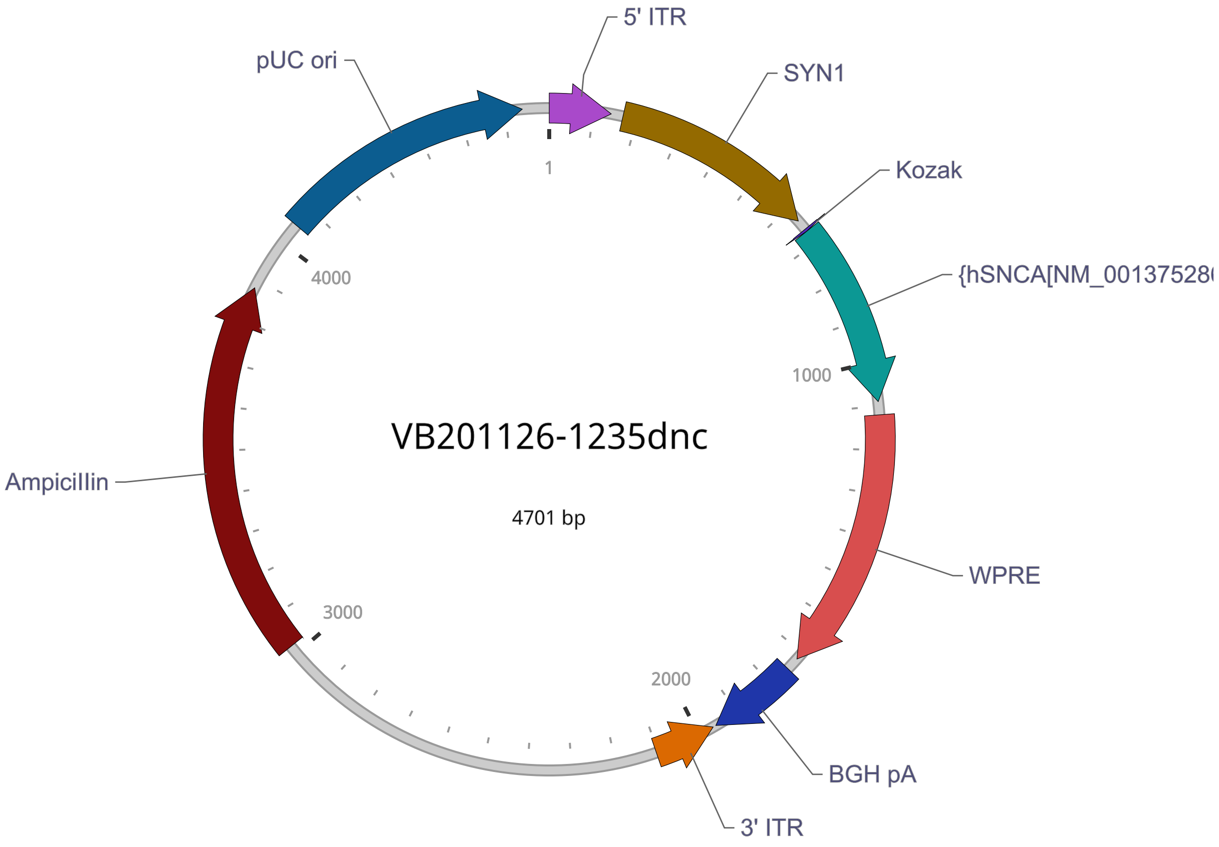


### Vector Components

| Name | Position | Size (bp) | Type | Description | Application notes |
|---|---|---|---|---|---|
| 5' ITR | 1-141 | 141 | ITR | AAV 5' inverted terminal repeat (functional equivalent of wild-type 5' ITR) | Allows replication of the viral genome and its packaging into virus. |
| SYN1 | 169-637 | 469 | Promoter | Human synapsin I promoter | Tissue specificity: Brain. Cell type specificity: Mature neurons. |
| Kozak | 662-667 | 6 | Miscellaneous | Kozak translation initiation sequence | Facilitates translation initiation of ATG start codon downstream of the Kozak sequence. |
| {hSNCA[NM_001375286.1]*(A53T)} | 668-1090 | 423 | ORF | *None* | *None* |
| WPRE | 1121-1718 | 598 | Miscellaneous | Woodchuck hepatitis virus posttranscriptional regulatory element | Enhances virus stability in packaging cells, leading to higher titer of packaged virus; enhances higher expression of transgenes. |
| BGH pA | 1749-1956 | 208 | PolyA_signal | Bovine growth hormone polyadenylation signal | Allows transcription termination and polyadenylation of mRNA transcribed by Pol II RNA polymerase. |
| 3' ITR | complement (1964-2104) | 141 | ITR | AAV 3' inverted terminal repeat | Allows replication of the viral genome and its packaging into virus. |
| AmpiciIIin | 3021-3881 | 861 | ORF | AmpiciIIin resistance gene | Allows E. coli to be resistant to ampiciIIin. |

| Name | Position | Size (bp) | Type | Description | Application notes |
|---|---|---|---|---|---|
| pUC ori | 4052-4640 | 589 | Rep_origin | pUC origin of replication | Facilitates plasmid replication in E. coli; regulates high-copy plasmid number (500-700). |

**Note:** Components added by user are listed in bold red text.

### Vector Sequence

```
   1 CCTGCAGGCA GCTGCGCGCT CGCTCGCTCA CTGAGGCCGC CCGGGCAAAG CCCGGGCGTC GGGCGACCTT TGGTCGCCCG
  81 GCCTCAGTGA GCGAGCGAGC GCGCAGAGAG GGAGTGGCCA ACTCCATCAC TAGGGGTTCC TTCTAGACAA CTTTGTATAG
 161 AAAAGTTGCT GCAGAGGGCC CTGCGTATGA GTGCAAGTGG GTTTTAGGAC CAGGATGAGG CGGGGTGGGG GTGCCTACCT
 241 GACGACCGAC CCCGACCCAC TGGACAAGCA CCCAACCCCC ATTCCCCAAA TTGCGCATCC CCTATCAGAG AGGGGGAGGG
 321 GAAACAGGAT GCGGCGAGGC GCGTGCGCAC TGCCAGCTTC AGCACCGCGG ACAGTGCCTT CGCCCCCGCC TGGCGGCGCG
 401 CGCCACCGCC GCCTCAGCAC TGAAGGCGCG CTGACGTCAC TCGCCGGTCC CCCGCAAACT CCCCTTCCCG GCCACCTTGG
 481 TCGCGTCCGC GCCGCCGCCG GCCCAGCCGG ACCGCACCAC GCGAGGCGCG AGATAGGGGG GCACGGGCGC GACCATCTGC
 561 GCTGCGGCGC CGGCGACTCA GCGCTGCCTC AGTCTGCGGT GGGCAGCGGA GGAGTCGTGT CGTGCCTGAG AGCGCAGCAA
 641 GTTTGTACAA AAAAGCAGGC TGCCACCATG GATGTATTCA TGAAAGGACT TTCAAAGGCC AAGGAGGGAG TTGTGGCTGC
 721 TGCTGAGAAA ACCAAACAGG GTGTGGCAGA AGCAGCAGGA AAGACAAAAG AGGGTGTTCT CTATGTAGGC TCCAAAACCA
 801 AGGAGGGAGT GGTGCATGGT GTGACAACAG TGGCTGAGAA GACCAAAGAG CAAGTGACAA ATGTTGGAGG AGCAGTGGTG
 881 ACGGGTGTGA CAGCAGTAGC CCAGAAGACA GTGGAGGGAG CAGGGAGCAT TGCAGCAGCC ACTGGCTTTG TCAAAAAGGA
 961 CCAGTTGGGC AAGAATGAAG AAGGAGCCCC ACAGGAAGGA ATTCTGGAAG ATATGCCTGT GGATCCTGAC AATGAGGCTT
1041 ATGAAATGCC TTCTGAGGAA GGGTATCAAG ACTACGAACC TGAAGCCTAA ACCCAGCTTT CTTGTACAAA GTGGGAATTC
1121 CGATAATCAA CCTCTGGATT ACAAAATTTG TGAAAGATTG ACTGGTATTC TTAACTATGT TGCTCCTTTT ACGCTATGTG
1201 GATACGCTGC TTTAATGCCT TTGTATCATG CTATTGCTTC CCGTATGGCT TTCATTTTCT CCTCCTTGTA TAAATCCTGG
1281 TTGCTGTCTC TTTATGAGGA GTTGTGGCCC GTTGTCAGGC AACGTGGCGT GGTGTGCACT GTGTTTGCTG ACGCAACCCC
1361 CACTGGTTGG GGCATTGCCA CCACCTGTCA GCTCCTTTCC GGGACTTTCG CTTTCCCCCT CCCTATTGCC ACGGCGGAAC
1441 TCATCGCCGC CTGCCTTGCC CGCTGCTGGA CAGGGGCTCG GCTGTTGGGC ACTGACAATT CCGTGGTGTT GTCGGGGAAG
1521 CTGACGTCCT TTCCATGGCT GCTCGCCTGT GTTGCCACCT GGATTCTGCG CGGGACGTCC TTCTGCTACG TCCCTTCGGC
1601 CCTCAATCCA GCGGACCTTC CTTCCCGCGG CCTGCTGCCG GCTCTGCGGC CTCTTCCGCG TCTTCGCCTT CGCCCTCAGA
1681 CGAGTCGGAT CTCCCTTTGG GCCGCCTCCC CGCATCGGGA ATTCCTAGAG CTCGCTGATC AGCCTCGACT GTGCCTTCTA
1761 GTTGCCAGCC ATCTGTTGTT TGCCCCTCCC CCGTGCCTTC CTTGACCCTG GAAGGTGCCA CTCCCACTGT CCTTTCCTAA
1841 TAAAATGAGG AAATTGCATC GCATTGTCTG AGTAGGTGTC ATTCTATTCT GGGGGGTGGG GTGGGGCAGG ACAGCAAGGG
1921 GGAGGATTGG GAAGAGAATA GCAGGCATGC TGGGGAGGGC CGCAGGAACC CCTAGTGATG GAGTTGGCCA CTCCCTCTCT
2001 GCGCGCTCGC TCGCTCACTG AGGCCGGGCG ACCAAAGGTC GCCCGACGCC CGGGCTTTGC CCGGGCGGCC TCAGTGAGCG
2081 AGCGAGCGCG CAGCTGCCTG CAGGGGCGCC TGATGCGGTA TTTTCTCCTT ACGCATCTGT GCGGTATTTC ACACCGCATA
2161 CGTCAAAGCA ACCATAGTAC GCGCCCTGTA GCGGCGCATT AAGCGCGGCG GGGGTGGTGG TTACGCGCAG CGTGACCGCT
2241 ACACTTGCCA GCGCCTTAGC GCCCGCTCCT TTCGCTTTCT TCCCTTCCTT TCTCGCCACG TTCGCCGGCT TTCCCCGTCA
2321 AGCTCTAAAT CGGGGGCTCC CTTTAGGGTT CCGATTTAGT GCTTTACGGC ACCTCGACCC CAAAAAACTT GATTTGGGTG
2401 ATGGTTCACG TAGTGGGCCA TCGCCCTGAT AGACGGTTTT TCGCCCTTTG ACGTTGGAGT CCACGTTCTT TAATAGTGGA
2481 CTCTTGTTCC AAACTGGAAC AACACTCAAC TCTATCTCGG GCTATTCTTT TGATTTATAA GGGATTTTGC CGATTTCGGT
2561 CTATTGGTTA AAAAATGAGC TGATTTAACA AAAATTTAAC GCGAATTTTA ACAAAATATT AACGTTTACA ATTTTATGGT
2641 GCACTCTCAG TACAATCTGC TCTGATGCCG CATAGTTAAG CCAGCCCCGA CACCCGCCAA CACCCGCTGA CGCGCCCTGA
2721 CGGGCTTGTC TGCTCCCGGC ATCCGCTTAC AGACAAGCTG TGACCGTCTC CGGGAGCTGC ATGTGTCAGA GGTTTTCACC
```

2801 GTCATCACCG AAACGCGCGA GACGAAAGGG CCTCGTGATA CGCCTATTTT TATAGGTTAA TGTCATGATA ATAATGGTTT
2881 CTTAGACGTC AGGTGGCACT TTTCGGGGAA ATGTGCGCGG AACCCCTATT TGTTTATTTT TCTAAATACA TTCAAATATG
2961 TATCCGCTCA TGAGACAATA ACCCTGATAA ATGCTTCAAT AATATTGAAA AAGGAAGAGT ATGAGTATTC AACATTTCCG
3041 TGTCGCCCTT ATTCCCTTTT TTGCGGCATT TTGCCTTCCT GTTTTTGCTC ACCCAGAAAC GCTGGTGAAA GTAAAAGATG
3121 CTGAAGATCA GTTGGGTGCA CGAGTGGGTT ACATCGAACT GGATCTCAAC AGCGGTAAGA TCCTTGAGAG TTTTCGCCCC
3201 GAAGAACGTT TTCCAATGAT GAGCACTTTT AAAGTTCTGC TATGTGGCGC GGTATTATCC CGTATTGACG CCGGGCAAGA
3281 GCAACTCGGT CGCCGCATAC ACTATTCTCA GAATGACTTG GTTGAGTACT CACCAGTCAC AGAAAAGCAT CTTACGGATG
3361 GCATGACAGT AAGAGAATTA TGCAGTGCTG CCATAACCAT GAGTGATAAC ACTGCGGCCA ACTTACTTCT GACAACGATC
3441 GGAGGACCGA AGGAGCTAAC CGCTTTTTTG CACAACATGG GGGATCATGT AACTCGCCTT GATCGTTGGG AACCGGAGCT
3521 GAATGAAGCC ATACCAAACG ACGAGCGTGA CACCACGATG CCTGTAGCAA TGGCAACAAC GTTGCGCAAA CTATTAACTG
3601 GCGAACTACT TACTCTAGCT TCCCGGCAAC AATTAATAGA CTGGATGGAG GCGGATAAAG TTGCAGGACC ACTTCTGCGC
3681 TCGGCCCTTC CGGCTGGCTG GTTTATTGCT GATAAATCTG GAGCCGGTGA GCGTGGAAGC CGCGGTATCA TTGCAGCACT
3761 GGGGCCAGAT GGTAAGCCCT CCCGTATCGT AGTTATCTAC ACGACGGGGA GTCAGGCAAC TATGGATGAA CGAAATAGAC
3841 AGATCGCTGA GATAGGTGCC TCACTGATTA AGCATTGGTA ACTGTCAGAC CAAGTTTACT CATATATACT TTAGATTGAT
3921 TTAAAACTTC ATTTTTAATT TAAAAGGATC TAGGTGAAGA TCCTTTTTGA TAATCTCATG ACCAAAATCC CTTAACGTGA
4001 GTTTTCGTTC CACTGAGCGT CAGACCCCGT AGAAAAGATC AAAGGATCTT CTTGAGATCC TTTTTTTCTG CGCGTAATCT
4081 GCTGCTTGCA AACAAAAAAA CCACCGCTAC CAGCGGTGGT TTGTTTGCCG GATCAAGAGC TACCAACTCT TTTTCCGAAG
4161 GTAACTGGCT TCAGCAGAGC GCAGATACCA AATACTGTTC TTCTAGTGTA GCCGTAGTTA GGCCACCACT TCAAGAACTC
4241 TGTAGCACCG CCTACATACC TCGCTCTGCT AATCCTGTTA CCAGTGGCTG CTGCCAGTGG CGATAAGTCG TGTCTTACCG
4321 GGTTGGACTC AAGACGATAG TTACCGGATA AGGCGCAGCG GTCGGGCTGA ACGGGGGGTT CGTGCACACA GCCCAGCTTG
4401 GAGCGAACGA CCTACACCGA ACTGAGATAC CTACAGCGTG AGCTATGAGA AAGCGCCACG CTTCCCGAAG GGAGAAAGGC
4481 GGACAGGTAT CCGGTAAGCG GCAGGGTCGG AACAGGAGAG CGCACGAGGG AGCTTCCAGG GGGAAACGCC TGGTATCTTT
4561 ATAGTCCTGT CGGGTTTCGC CACCTCTGAC TTGAGCGTCG ATTTTTGTGA TGCTCGTCAG GGGGGCGGAG CCTATGGAAA
4641 AACGCCAGCA ACGCGGCCTT TTTACGGTTC CTGGCCTTTT GCTGGCCTTT TGCTCACATG T

Deleted from original sequence:

▼ 824-1247

G

### Validation by Restriction Enzyme Digestion

| Restriction Enzymes | Cutting Sites | DNA Fragments (bp) |
|---|---|---|
| NcoI | 667, 1534 | 867, 3834 |
| ApaLI | 1335, 2640, 3137, 4383 | 1305, 497, 1246, 1653 |
| DraIII | 2413 | 4701 |
| ApaLI+NcoI | 667, 1335, 1534, 2640, 3137, 4383 | 668, 199, 1106, 497, 1246, 985 |
| ApaLI+DraIII | 1335, 2413, 2640, 3137, 4383 | 1078, 227, 497, 1246, 1653 |
